# UVREK: Development and analysis of an expression profile knowledgebase of biomolecules induced by ultraviolet radiation exposure

**DOI:** 10.1101/2024.03.31.587452

**Authors:** Shanmuga Priya Baskaran, Janani Ravichandran, Priya Shree, Vinayak Thengumthottathil, Bagavathy Shanmugam Karthikeyan, Areejit Samal

## Abstract

Humans encounter diverse environmental factors which can have impact on their health. One such environmental factor is the ultraviolet (UV) radiation, which is part of the physical component of the exposome. UV radiation is the leading cause of skin cancer and is a significant global health concern. A large body of published research has been conducted to uncover the mechanisms underlying the adverse outcomes of UV radiation exposure on living beings. These studies involve identifying the biomolecules induced upon UV radiation exposure. A few previous efforts have attempted to compile this information in the form of a database, but such earlier efforts have certain limitations. To fill this gap, we present a structured database named UVREK, containing manually curated data on biomolecules induced by UV radiation exposure from published literature. UVREK has compiled information on 985 genes, 470 proteins, 54 metabolites and 77 miRNAs along with their metadata. Thereafter, an enrichment analysis performed on the human gene set of UVREK database showed the importance of transcription related processes in UV related response, and enrichment of pathways involved in cancer and aging. While significantly contributing towards characterizing the physical component of the exposome, we expect the compiled data in UVREK will serve as a valuable resource for development of better UV protection mechanisms such as UV sensors and sunscreens. Noteworthy, UVREK is the only resource to date, compiling varied types of biomolecular responses to UV radiation with corresponding metadata. UVREK is openly accessible at: https://cb.imsc.res.in/uvrek/.

## 1. Introduction

Humans are constantly exposed to myriad stressors in the environment that have impact on biological processes and eventually health. Such environmental exposures are referred to as ‘exposome’, which encapsulates the entire spectrum of environmental influences encountered by individuals throughout their lifetime. Specifically, human exposome encompasses chemical, physical, biological and social factors, which play a significant role in shaping our health and well-being [1–6]. In terms of understanding the impact of exposome on health, significant research effort has been devoted towards characterizing the chemical component of the exposome. These research efforts on chemical exposome include the Blood Exposome [7,8], CTD [9], DEDuCT [10,11], ExHuMId [12], Exposome-Explorer [13,14,2], FCCP [15], Human Indoor Exposome Database [16], NeurotoxKb [17], TExAs [18], T3DB [19,20], ViCEKb [21] among others. In contrast to the chemical exposome, much less effort has been devoted towards characterizing the physical component of the exposome and its impact on human health.

Physical components (or stressors) in the exposome include heat, noise and radiation, which can play a substantial role in shaping human health. Among these exposures, ultraviolet (UV) radiation constitutes a significant portion of everyday human exposure and exerts both harmful and beneficial effects [22–25]. UV radiation is a form of non-ionizing radiation with a wavelength ranging from 100 nm to 400 nm, and accounts for 5% of the total solar spectrum [26,27]. Based on the wavelength, UV radiation is categorized into three groups namely, UVC (100–280 nm), UVB (280–315 nm) and UVA (315–400 nm). Note that the longer the wavelength, the deeper it can penetrate into the skin layers. In case of UV, UVA has the longest wavelength and is capable of reaching the dermis of the skin. UVB mostly affects the epidermis and is capable of reaching the papillary dermis of skin [24,27]. UVC radiation, which has the shortest wavelength among the UV types and is known for its germicidal effects [28], gets filtered by atmospheric ozone and does not reach the skin in natural conditions.

UV radiation exposure is a major environmental factor with impact on skin health and contributes significantly to skin aging. It also induces DNA damage, oxidative stress, free radical production, cell proliferation and inflammation, photoreactions and immunosuppression, which are all linked to tumorigenesis [24,29,30]. UV radiation also increases the risk of skin cancer and cataracts, and there is also evidence indicating that both UVB and UVA promote melanoma development [24,31]. Apart from the documented negative effects, UV radiation is also known to have positive effects. UV radiation is well-known to induce vitamin D production in the skin. It has also been reported to improve skin related conditions such as psoriasis, vitiligo and localized scleroderma [32]. UV radiation induces nitric oxide production which in turn has effects such as blood pressure reduction [33], stimulation of pigmentation [34] and antimicrobial activity [35]. Due to these reasons, it is crucial to understand the biomolecular responses to UV radiation exposure, which in turn will enable the appropriate use of UV radiation for benefits or seek proper protection from unwarranted exposure [36,26].

As the information on biomolecules induced by UV radiation exposure from previous research is scattered across published literature, there is a need to compile such information in a structured database centered on UV radiation exposure and their subsequent biomolecular responses. In this direction, there have been few efforts towards the development of radiation-specific databases, such as UVGD 1.0 [26], RadAtlas 1.0 [36] and the Radiation Genes database [37]. However, RadAtlas 1.0 and the Radiation Genes database have compiled data related to ionizing radiation, while UVGD 1.0 does not compile information on the sampling time, dose, dosage rate, and other experimental details, and is also limited to gene-related data (**Table 1**). Addressing these gaps, by including details such as sampling time, dose, dosage rate, and compilation of other experimental conditions under which a particular biomolecule was induced, is crucial for enhancing the utility of such a database from the perspective of future applications [38]. Additionally, compiling datasets pertaining to diverse biomolecules such as gene, protein, metabolite, and miRNA expression profiles will provide a sophisticated platform for advancing our understanding of the biological impact of UV radiation exposure on human physiology.

**Table 1:** Comparison of the features in UVREK database with two previously published resources namely, UVGD 1.0 [26] and RadAtlas 1.0 [36].

| Feature | UVREK | UVGD 1.0 | RadAtlas 1.0 |
| --- | --- | --- | --- |
| Radiation type used | Yes | No | No |
| Dosage rate | Yes | No | No |
| Dose | Yes | No | No |
| Sampling time | Yes | No | No |
| Tested organisms | Human<br>Mouse<br>Rat | Human | Human |
| Tested cell/tissue type | Yes | No | No |
| Expression profile of genes | Yes | Yes | Yes |
| Number of compiled genes | 985 | 663 | 598 |
| Expression profile of proteins | Yes | No | No |
| Expression profile of metabolites | Yes | No | No |
| Expression profile of miRNAs | Yes | No | No |

In this pursuit of establishing a UV-centric database, we systematically compiled and curated data on biomolecules induced by UV radiation exposure from 320 filtered published research articles, focusing specifically on studies involving humans and rodents. Specifically, we have compiled biomolecule response data obtained in normal, healthy cells or tissues exposed to UV radiation. This choice was influenced by the central motivation to contribute towards characterizing the physical component of the exposome, and including specimens of diseased state will distort this purpose [1,6]. Importantly, capturing responses in normal cells combined with dose information will help in the detection of potential biomarkers, which in turn can help in accessing the exposure level of an individual and help in risk stratification [39,40,4,41]. From the filtered articles, we meticulously collected biomolecular responses induced by UV radiation, including gene, protein, metabolite, and miRNA expression profiles, and mapped them to unique identifiers. This systematic approach resulted in a comprehensive compilation of biomolecule expression profiles, encompassing 985 genes, 470 proteins, 54 metabolites, and 77 miRNAs, sourced from studies spanning diverse cell types, tissue types, and experimental conditions. Subsequently, in order to widely share this curated dataset, we created an online resource, Ultraviolet Radiation Expression Knowledgebase (UVREK), which is openly accessible at: https://cb.imsc.res.in/uvrek. Furthermore, to elucidate the molecular mechanisms and biological processes associated with UV radiation exposure, we performed extensive analyses such as GO term enrichment, pathway enrichment, and protein-protein interaction (PPI) network construction by leveraging the compiled dataset in UVREK. These analyses yielded valuable insights into the biological effects of UV radiation and the intricate crosstalk between pathways involved in the response to UV exposure, thereby enhancing our understanding of UV-induced biomolecular changes.

## 2. Methods

### 2.1 Compilation of published research articles relevant to UV radiation exposure

The primary objective of this study is to compile expression profiles of biomolecules such as genes, proteins, metabolites and miRNAs induced by Ultraviolet (UV) radiation exposure. To this end, we conducted a comprehensive literature search on PubMed (https://pubmed.ncbi.nlm.nih.gov/) by using the following structured query (which includes related keywords): ‘(human[ORGN] OR “Homo sapiens”[ORGN] OR rat[ORGN] OR “Rattus norvegicus”[ORGN] OR mouse[ORGN] OR “Mus musculus”[ORGN] OR mice[ORGN] OR murine[ORGN]) AND ((UV*[TIAB] OR ultraviolet*[TIAB]) AND (radiation[TIAB] OR irradiation[TIAB] OR rays[TIAB])) AND ((gene*[TIAB] OR genome[TIAB] OR genomic*[TIAB] OR microarray[TIAB] OR “gene expression”[TIAB] OR gene-expression[TIAB]) OR (protein*[TIAB] OR proteome[TIAB] OR proteomic*[TIAB] OR “gene-product”[TIAB] OR “gene product”[TIAB] OR “protein expression”[TIAB] OR “protein-expression”[TIAB]) OR (metabolite*[TIAB] OR metabolome[TIAB] OR metabolomic*[TIAB] OR metabolic[TIAB] OR metabolize[TIAB] OR metabolise[TIAB])) AND (effect[TIAB] OR exposure[TIAB] OR response[TIAB] OR injury[TIAB] OR biomarker[TIAB] OR signature[TIAB])’

The above query search was last performed on 12 February 2022 and yielded a total of 5553 published research articles (**Figure 1**). Thereafter, these articles were screened based on specific inclusion and exclusion criteria (**Figure 1**), which are as follows:

i. Experiments reported in the article should have utilized UV radiation;
ii. Experiments should have been conducted on humans or rodents;
iii. Experiments should have been conducted on normal cells or tissues, excluding cancer cells, gene knockout studies or any conditions involving disease;
iv. Proper controls, without UV irradiation, must be present;
v. The article should report statistically significant change (p-value ≤ 0.05) in the expression profiles of genes or proteins or metabolites or miRNAs;
vi. Articles reporting results from mixed effects, such as a combined effect of UV radiation and chemical exposure, were excluded;
vii. Meta-analysis studies were excluded;
viii. Review articles, articles in languages other than English, and articles inaccessible were excluded.

**Figure 1:**
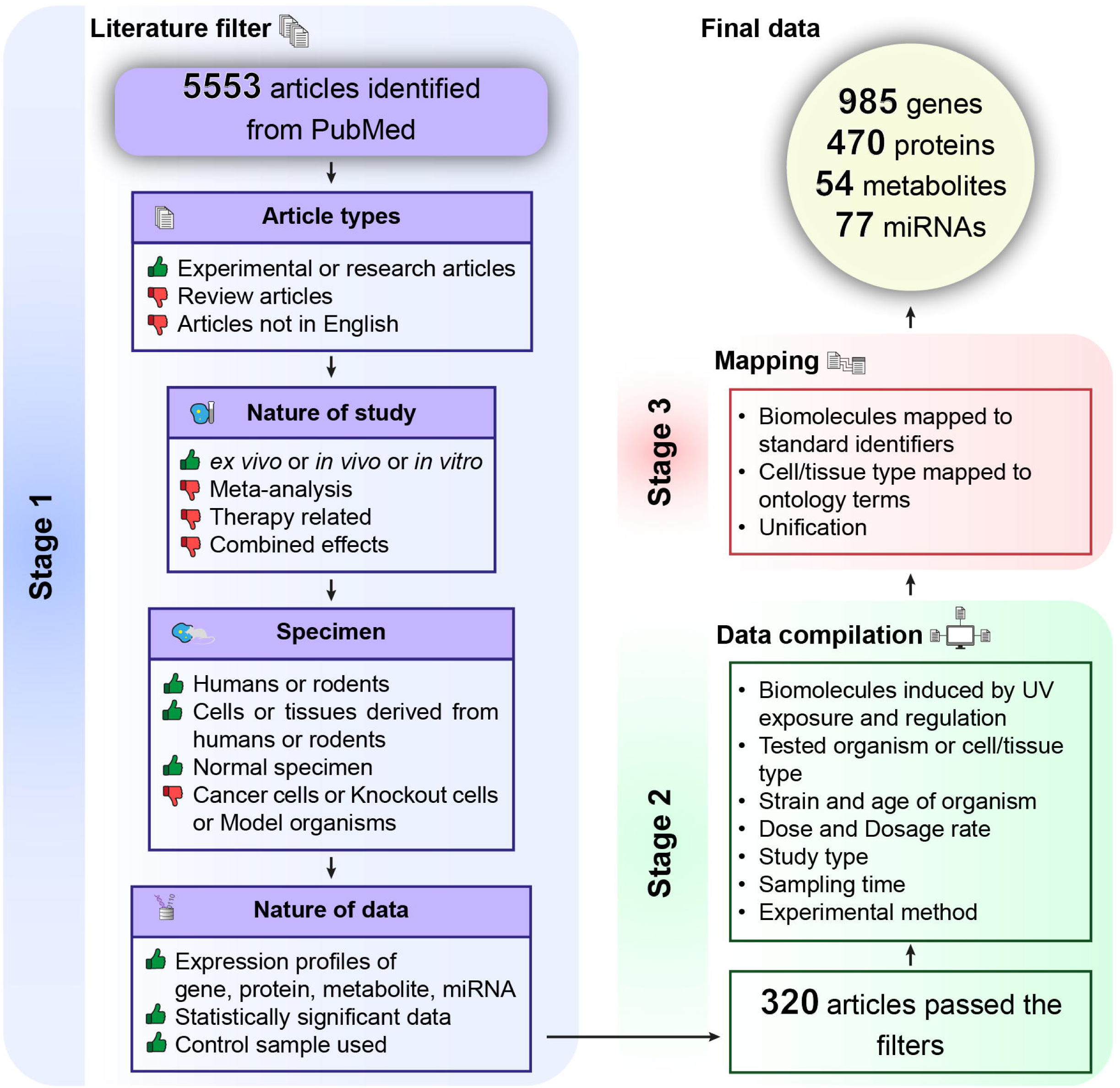
A systematic workflow for identifying published research articles containing information on the expression profiles of genes, proteins, metabolites, and miRNAs induced upon exposure to Ultraviolet (UV) radiation in humans and rodents.

Altogether, this screening workflow led to a subset of 320 PubMed Identifiers (PMIDs) or published research articles, which were shortlisted for the next stage (**Figure 1**).

### 2.2 Data compilation and annotation

From the 320 shortlisted research articles relevant to UV radiation exposure, an exhaustive manual extraction of the expression profiles for genes, proteins, metabolites and miRNAs was performed. Simultaneously, metadata associated with the expression profiles was also meticulously collected. The metadata includes details such as the organism under study or organism from which the tested cell or tissue was derived, the strain and age of the organism, the cell or tissue from which the data was obtained, the type of UV radiation used for the experiment, the dose and dosage rate of radiation, the study type (*in vivo* or *in vitro* or *ex vivo*), sampling time, type of omics data collected, and the experimental method used for data acquisition. Noteworthy, some published studies have simply reported that UV radiation has been utilized and have not indicated the specific UV type used. In such instances, we have designated the type of UV radiation as ‘UV radiation-General’.

Subsequent to the data extraction, names of genes, proteins, metabolites and miRNAs were mapped to identifiers (IDs) from standard databases. This process involved:

i. Mapping of a Gene to GeneSymbol, Organism-specific Database ID, Official name, Ensembl ID [42], NCBI Gene ID (https://www.ncbi.nlm.nih.gov/gene/), and corresponding organism information from NCBI database.
ii. Mapping of a Protein to UniProt Name, UniProt ID, Gene Symbol, Organism-specific Database ID, Official Name, and corresponding organism information from UniProt database [43].
iii. Mapping of a Metabolite to KEGG ID, Official Name, synonyms from KEGG database [44,45] (https://www.genome.jp/kegg/), and to CAS ID, PubChem ID and ChEBI ID using CAS (https://www.cas.org/), PubChem [46] and ChEBI [47] databases, respectively.
iv. Mapping of a miRNA to Accession code, Symbol, Description, Mature sequence, Gene family and corresponding organism information using miRBase [48].

Biomolecules (i.e. genes or proteins or metabolites or miRNAs) which could not be mapped to standard identifiers were excluded. Additionally, the tested cell or tissue type corresponding to biomolecule data was mapped to the standard identifiers using Ontology Lookup Service (OLS) (https://www.ebi.ac.uk/ols4/). Information on dose, dosage rate and sampling time was standardized and unified to units of J/m² for dose, W/m² for dosage rate, and hours for sampling time, respectively.

Finally, we compiled a dataset of expression profiles of biomolecules relevant to UV radiation exposure comprising of 985 genes (with Gene Symbols), 470 proteins (with UniProt IDs), 54 metabolites (with CAS IDs) and 77 miRNAs (with Accession codes) (**Supplementary Tables S1-S4**). Note that a given gene, protein, metabolite or miRNA in our compiled dataset may have been reported in multiple studies spanning different organisms, cell types, tissue types or experimental conditions. Therefore, our compiled dataset comprises 2297 gene expression profiles, 1228 protein expression profiles, 209 metabolite expression profiles and 162 miRNA expression profiles (**Supplementary Tables S1-S4**). Notably, 811 out of the 985 genes in our compiled dataset belong to humans and have unique Gene Symbols, and moreover, this set of 811 genes (**Supplementary Table S5**) was used as the input dataset for subsequent analysis, unless explicitly stated otherwise.

### 2.3 Web interface and database management system

Leveraging the compiled information, we have developed an online database with a user-friendly web interface namely, **U**ltra**V**iolet **R**adiation **E**xpression **K**nowledgebase (UVREK). UVREK compiles information on four types of biomolecules, namely, genes, proteins, metabolites and miRNAs, induced by UV radiation exposure. Additionally, UVREK provides mapping of the biomolecule information to standard databases and includes associated metadata. UVREK is openly accessible for academic research at: https://cb.imsc.res.in/uvrek/.

Within the web interface of UVREK, we employed MariaDB (https://mariadb.org/) to store the compiled dataset and utilized Structured Query Language (SQL) for data retrieval. The web interface was built using PHP (https://www.php.net/) along with customized HTML, CSS, jQuery (https://jquery.com/), and Bootstrap 5 (https://getbootstrap.com/docs/5.0/). The UVREK database is hosted on an Apache webserver (https://httpd.apache.org/) running on Debian 9.4 Linux Operating System.

### 2.4 GO term enrichment analysis

We conducted GO term enrichment analysis to elucidate the molecular mechanisms, biological processes and pathways associated with gene responses to UV radiation exposure. This analysis was performed for the set of 811 human genes using the Enrichr webserver [49] and the results were obtained for GO terms across three ontologies: biological process, cellular component, and molecular function. For each of the three ontologies, we extracted the significantly enriched (p-value ≤ 0.01) GO terms along with the corresponding number of genes associated with UV radiation exposure for each term. Subsequently, the extracted GO terms were sorted based on the number of associated genes, and the top 10 enriched terms (**Supplementary Table S6**) were visualized using the Matplotlib library (https://matplotlib.org/).

### 2.5 Pathway enrichment analysis

We also performed pathway enrichment analysis for the set of 811 human genes induced by UV radiation exposure. For this, we utilized the ‘KEGG 2021 Human’ gene set library of Enrichr [49] for pathway associations. This analysis yielded a ranked list of pathways wherein the ranking was based on the p-value quantifying the similarity of the input gene set with the pathway gene set. Further we filtered the KEGG pathways with a gene set size (i.e., number of genes associated with UV exposure response) greater than 25, aiming to mitigate overestimation of resultant statistics. Additionally, we applied a p-value cut-off of ≤ 0.01, resulting in the identification of 53 enriched KEGG pathways (**Supplementary Table S7**) within the input gene set. Moreover, we manually mapped each of these 53 enriched pathways to their corresponding hierarchical classifications, as provided by the KEGG BRITE database (https://www.genome.jp/kegg/brite.html) (**Supplementary Table S7**). The enriched pathways in the input gene set were visualized using bubble plots generated utilizing Matplotlib library (https://matplotlib.org/).

For a broader understanding of the relationships between the 53 enriched pathways, we constructed a pathway similarity network by computing the Jaccard index between the enriched pathways based on their gene sets. Within this network, two pathways were linked by an edge if their respective Jaccard index exceeded 0.2, a threshold delineating the top quartile of the Jaccard index distribution. Visualization of the pathway similarity network was performed using Cytoscape version 3.9.1 [50].

### 2.6 Protein interaction network based analysis

To explore the functional relationships among the 811 human genes induced by UV radiation exposure, we leveraged the NetworkAnalyst 3.0 webserver (https://www.networkanalyst.ca/) to construct the associated protein-protein interaction (PPI) network [51]. For this construction, we employed the STRING database [52] within the NetworkAnalyst webserver, and moreover, set the confidence score threshold to be greater than 0.9. Of note, only interactions with known experimental evidence were considered. Thereafter, we derived a first-order network, filtered the nodes for a degree of 5, and visualized the largest connected component of the first-order network (**Supplementary Tables S8-S9**) using Gephi 0.9.7 [53].

In this analysis, we further categorized the standardized terms corresponding to cells or tissues (wherein expression profiles were obtained) into broader classifications, encompassing the following categories: Skin, Eye, Blood, Connective tissue, Liver, Intestine, Thyroid gland, Ear, Testis, Spleen, and Others (**Supplementary Tables S1-S4**). Based on this categorization, we observed that among the compiled data in UVREK, majority of the gene expression profiles were obtained in skin-specific tissues or cells. Therefore, we also constructed another PPI network which is skin-specific by incorporating genes only in the skin category induced by UV radiation exposure. While constructing the skin-specific PPI network within the NetworkAnalyst webserver, we additionally filtered the skin-specific nodes using the information from Genotype-Tissue Expression (GTEx) portal [54]. Subsequently, we derived a first-order network, filtered the nodes for a degree of 5, and visualized the largest connected component of the first-order network (**Supplementary Tables S10-S11**) using Gephi 0.9.7 [53].

## 3. Results and Discussion

### 3.1 UVREK: Ultraviolet Radiation Expression Knowledgebase

Humans are exposed to various forms of radiation, with UV radiation constituting a significant portion of everyday exposure. Our primary objective is to establish a comprehensive database focused on UV radiation exposure and the corresponding biomolecular response, encompassing gene, protein, metabolite, and miRNA expression profiles. By adhering to a structured workflow (**Figure 1**; **Methods**), we filtered a curated set of 320 published research articles containing pertinent information on biomolecular responses triggered by UV radiation exposure across transcriptomic, proteomic, metabolomic, and miRNA expression studies. We collected the expression profiles of genes, proteins, metabolites, and miRNAs from the filtered research articles along with the corresponding metadata such as the experimental conditions, including details on tested organisms, organ or tissue or cell types, and dosage information (**Figure 1**; **Methods**). Finally, we annotated and compiled a non-redundant set of 985 genes, 470 proteins, 54 metabolites, and 77 miRNAs that have been shown to significantly respond to UV radiation exposure in published studies. Using this compiled dataset, we built UVREK, a dedicated resource on UV radiation induced biomolecular expression profiles, which is openly accessible for academic research at: https://cb.imsc.res.in/uvrek/.

**Figure 2** visualizes the connections between data and metadata compiled in UVREK. That is, the connections among various specifications of UV radiation exposures and biomolecular responses across the compiled data within UVREK. Moreover, **Figure 2** provides a comprehensive view of the nature of the filtered studies by displaying the interconnections between experimental conditions, specimen, and expression profiles. **Figure 3** displays screenshots of different pages and navigation options on the UVREK website, showcasing the structured online platform and user-friendly web interface of the database designed to ensure easy navigation and convenient access to the aggregated data for users (**Figure 3a**). The ‘BROWSE’ tab enables users to explore content in two ways: (i) Browse by UV type, or (ii) Browse by Biomolecules option (**Figure 3b**). Browse by UV type enables users to explore data corresponding to various UV types: UVA, UVB, UVC, UVA+UVB and UVR-General (UV radiation-General). Similarly, Browse by Biomolecules enables users to explore UV radiation exposure induced biomolecule data on genes, proteins, metabolites, and miRNAs individually. **Figure 3c** displays the tabular output from UVREK for genes induced by UV. **Figure 3c** also displays a column in the tabular output which contains a link for the experimental information associated with the expression profile, facilitating access to all the metadata related to the corresponding data (**Figure 3d**). In UVREK, clicking on the standardized identifiers (IDs) of the biomolecules will provide detailed information in a consolidated tabular format on all relevant literature and experimental conditions in which the biomolecules have been studied (**Figure 3e**). Furthermore, the ‘SEARCH’ tab allows users to query the UVREK database for biomolecules in two ways: (i) Search by ID, or (ii) Search by UV details. Search by ID enables users to search for the specific biomolecules using the corresponding identifiers or name (**Figure 3f**). Search by UV details allows users to search with details such as the UV radiation type, dose and dosage rate, and the results are presented in a tabulated format (**Figure 3g**). Additionally, users can access data organized based on PMID through the ‘PMID’ tab under the ‘SEARCH’ tab.

**Figure 2:**
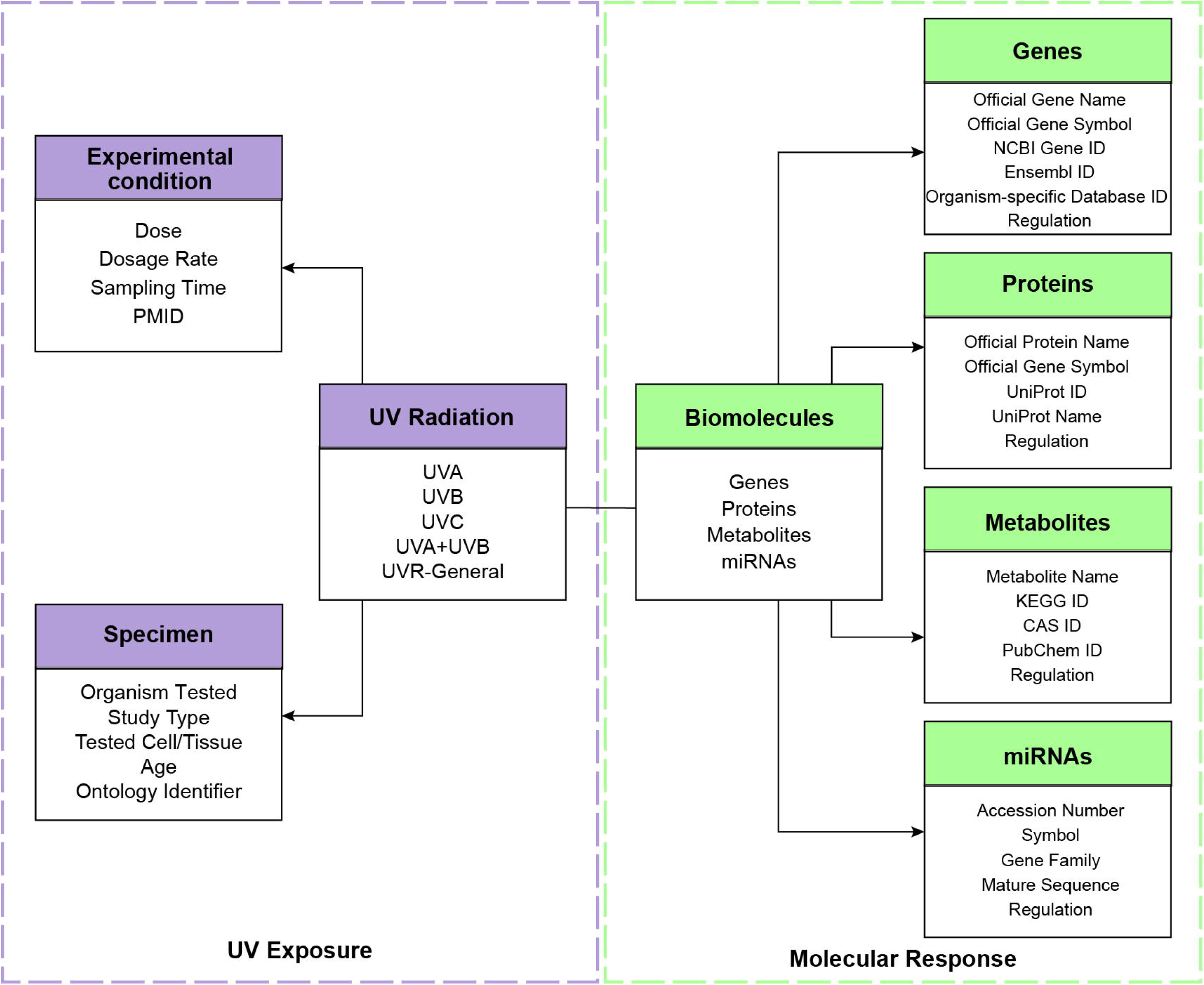
An illustrative schematic depicting the connections between data and metadata. This figure showcases the interconnection between various specifications of UV radiation exposure and biomolecular responses. UV radiation is connected to experimental conditions and organisms in which experiments were conducted, as well as the study type, while biomolecules such as genes, proteins, metabolites, and miRNAs are linked to their mapping to unique identifiers in standard databases.

**Figure 3:**
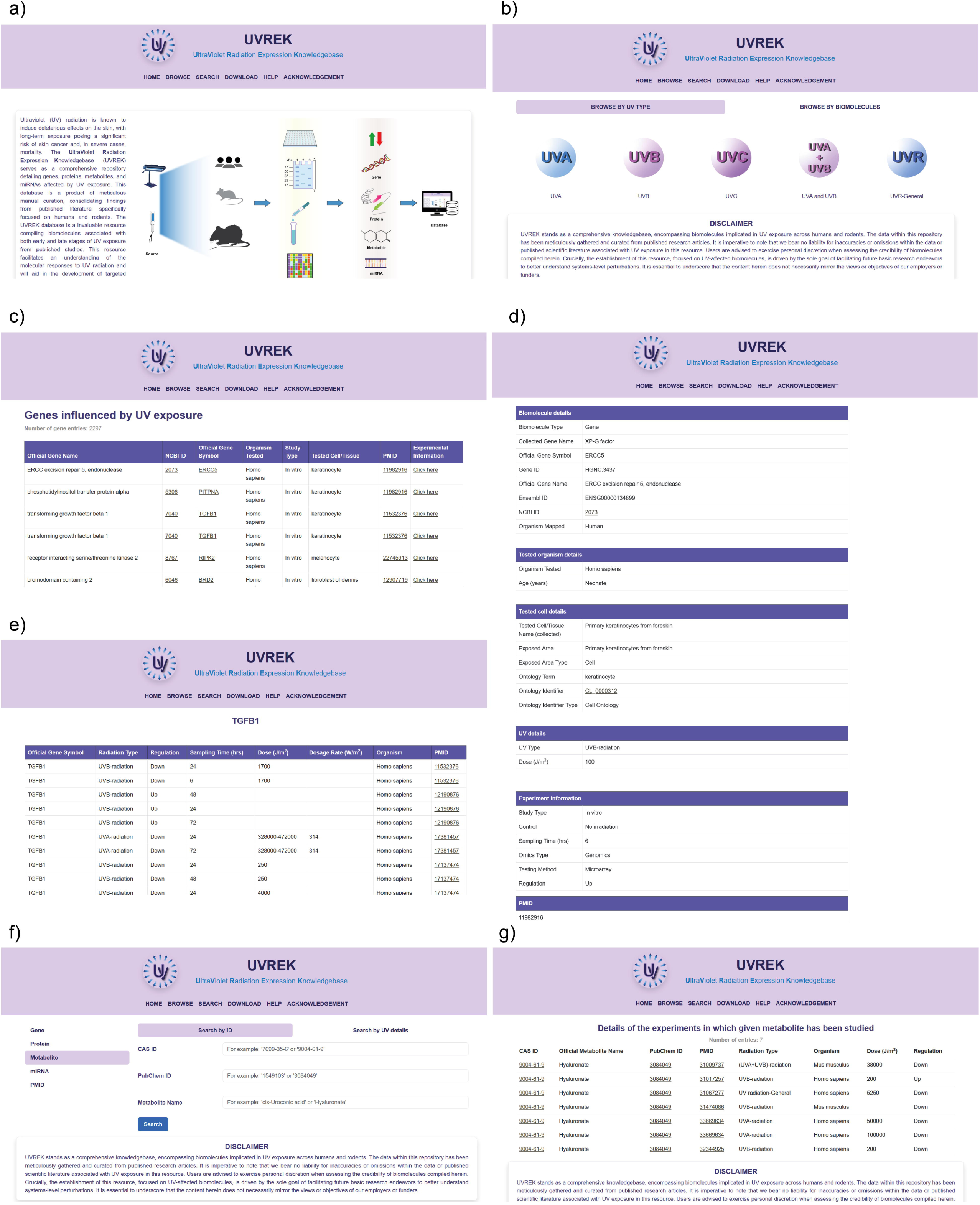
The screenshots illustrate the user interface and navigation features of the UVREK website. **(a)** Screenshot of the UVREK homepage which has a structured layout for ease of navigation and access to compiled data. **(b)** BROWSE tab showing two primary ways of exploring the content: Browse by UV type or Browse by Biomolecules. **(c)** BROWSE tab enables users to explore UV radiation exposure induced biomolecule data on genes, proteins, metabolites, and miRNAs, with results conveniently displayed in tabular formats. Last column of the tabular output has links to the associated experimental information, and clicking the corresponding link will lead to the page shown in part (d). **(d)** Screenshot of the page containing detailed information on the experimental conditions and other metadata associated with the expression profile of the corresponding biomolecule. **(e)** Screenshot of the tabular column obtained as a result of clicking on the standardized identifiers of a biomolecule. The resulting page displays information on all studies within UVREK in which the biomolecule has been reported. **(f)** Screenshot of the SEARCH page enabling users to query the compiled biomolecules in UVREK database based on their identifiers or UV-specific details, or PMIDs. **(g)** Screenshot of the results obtained in a tabular format from an example query search.

### 3.2 FAIR principles compliance

The UVREK database is compliant with the Findable (F), Accessible (A), Interoperable (I), and Reusable (R) data principles (FAIR) [55] which is evident from the following:

a. It has a stable, permanent web address (F, A).
b. Entries are uniquely identified by stable URLs (F).
c. Biomolecules are identified by unique IDs from standardized databases, and corresponding links are provided as detailed in the **Methods** section (F, I).
d. References are identified by PMIDs (F, R).
e. Tested cell or tissue types are identified by unique ontology identifiers (F).
f. The database does not require the creation of an account for utilization (A).
g. A HELP page is available providing guidance on how to use the database (A).
h. Data and metadata are available in a human-readable format, i.e., HTML (A), and downloadable in a machine-readable format, i.e., CSV (A, R).
i. The license (CC BY-NC 4.0) for use of data compiled in UVREK is available (R).

It is important to note that UVREK is hosted in the institutional data repository, ensuring long-term sustainability.

### 3.3 UVREK database statistics and comparison with other resources

**Table 1** presents a comparison of the features of UVREK with two previously published UV radiation specific resources namely, UVGD 1.0 [26] and RadAtlas 1.0 [36]. Noteworthy, UVREK has compiled more detailed experimental information, including sampling time, dose, dosage rate, and data specific to human, mouse, and rat. In comparison, UVGD 1.0 [26] and RadAtlas 1.0 [36] do not compile such detailed experimental information. Moreover, the type of UV radiation employed in the published experiments has been systematically documented in UVREK. Notably, UVREK also compiles expression profiles for proteins, metabolites and miRNAs, a feature absent in UVGD 1.0 [26] and RadAtlas 1.0 [36].

**Figure 4a** summarizes the number of unique biomolecules namely, genes, proteins, metabolites and miRNAs, reported to have been induced by UV exposure in the different tested organisms namely, *Homo sapiens* (human), *Mus musculus* (mouse), *Rattus norvegicus* (rat), in the published studies compiled in UVREK. This plot reveals that, in published studies on humans, genes predominate followed by proteins, while a similar trend is also seen for studies in mouse. In contrast, fewer genes have been reported in rats compared to proteins in the published studies curated in UVREK (**Figure 4a**).

**Figure 4:**
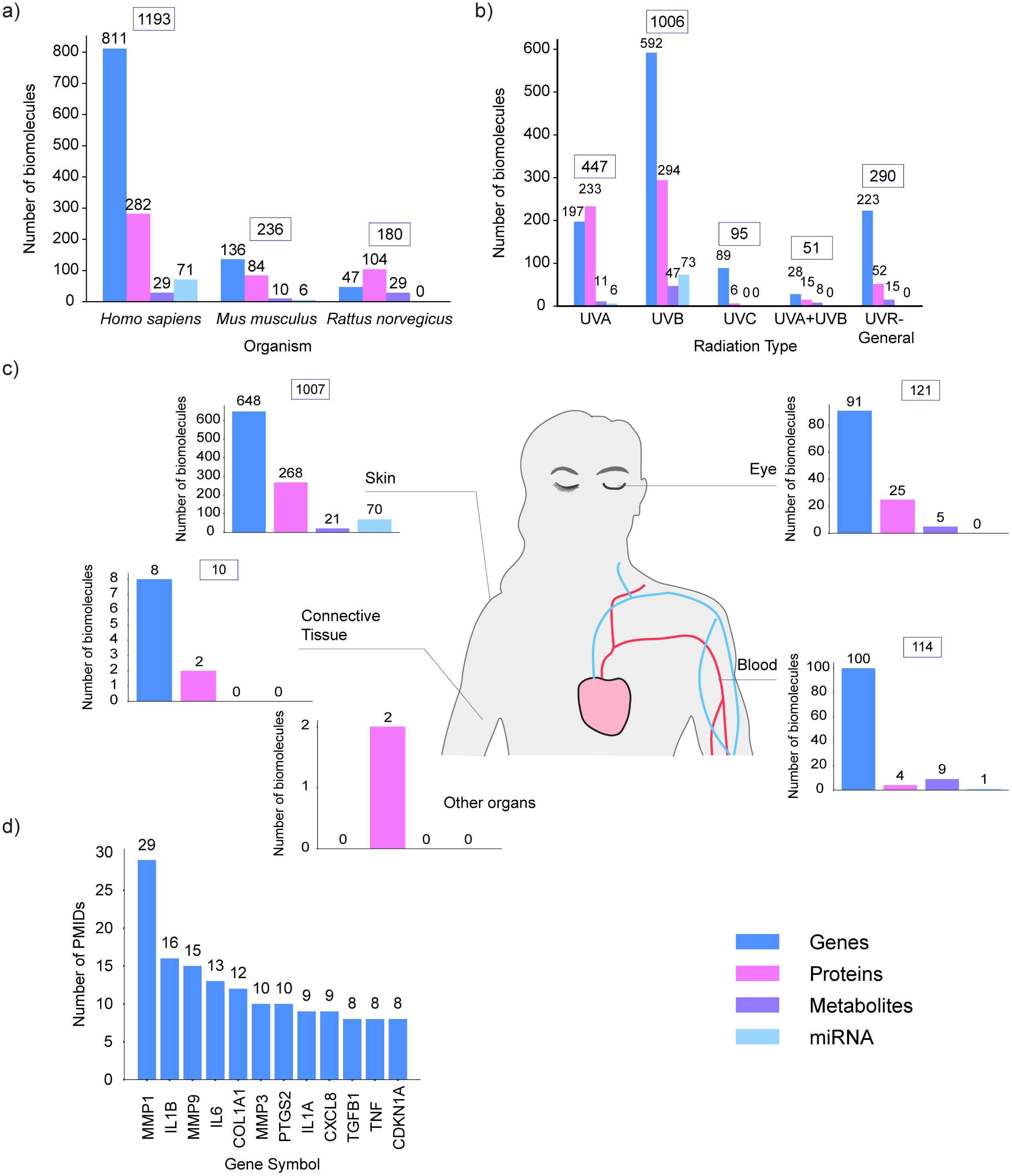
Distribution of biomolecules induced by UV radiation exposure based on various criteria. **(a)** Distribution of biomolecules based on the tested organism namely, *Homo sapiens* (human), *Mus musculus* (mouse), or *Rattus norvegicus* (rat). **(b)** Distribution of biomolecules based on the type of UV radiation employed. **(c)** Distribution of biomolecules specific to humans based on the exposed organ or tissue types. **(d)** The top 10 genes based on the number of associated published studies or PubMed Identifiers (PMIDs) compiled in UVREK. The displayed distributions in this figure were obtained based on unique identifiers for genes, proteins, metabolites and miRNAs in UVREK as described in the Methods section.

**Figure 4b** displays the number of biomolecules induced by different UV radiation types based on compiled data from published studies in UVREK. In terms of the number of genes reported in published studies, UVB is the highest with 592 genes followed by the UV radiation-General category with 223 genes. Despite UVA being the predominant UV radiation type both in terms of skin penetration and fraction of UV in the atmosphere [27,56], UVB has higher number of associated biomolecules in UVREK.

Further, based on the broader classification of cell or tissue type in the compiled experimental studies in UVREK (**Methods**), we analyzed the expression profiles of biomolecules specific to human as illustrated in **Figure 4c**. The number of biomolecules reported in published studies compiled in UVREK and specific to skin is highest (1007 biomolecules of which 648 are genes), followed by eye (121 biomolecules of which 91 are genes), blood (114 biomolecules of which 100 are genes) and connective tissue (10 biomolecules of which 8 are genes). Notably, within the compiled studies in UVREK, tissues such as liver, intestine, thyroid gland, ear, testis and spleen have been studied in mouse and rat, but not in humans (**Supplementary Tables S1-S4**).

In UVREK, we compiled 811 human genes which have been reported to be induced by UV radiation exposure across the published studies. We next analyzed the number of published studies (across the 320 PMIDs compiled in UVREK) that implicate each of these 811 human genes (**Supplementary Table S5**). **Figure 4d** displays the top 10 genes based on the number of published studies compiled in UVREK associated with each gene. Specifically, Matrix Metallopeptidase 1 (MMP1) was reported in maximum number of studies (29 PMIDs), followed by Interleukin 1 beta (IL1B) and Matrix Metallopeptidase 9 (MMP9).

### 3.4 Detailed analysis of genes induced by UV radiation exposure

GO enrichment analysis provides valuable insights on complex biological processes associated with a set of biomolecules (e.g. genes). Hence, we performed GO enrichment analysis for the 811 human genes in UVREK that are induced by UV radiation exposure. In particular, we performed this enrichment analysis for the three different ontologies: biological process, cellular component, and molecular function (**Methods**), and **Figure 5** displays the top 10 enriched GO terms for each of these ontologies (**Supplementary Table S6**).

**Figure 5:**
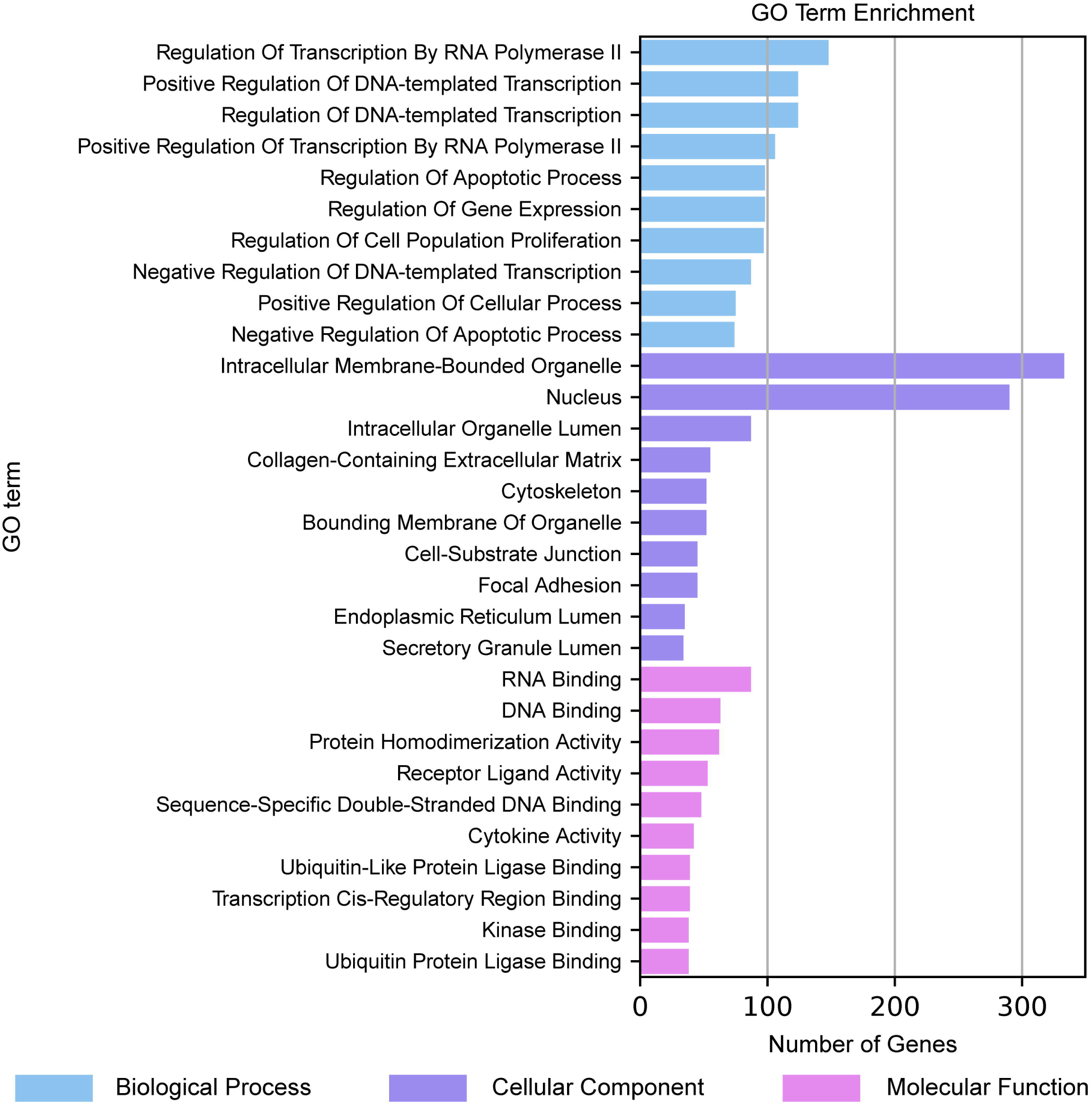
Bar chart displaying the top 10 enriched gene ontology (GO) terms across three categories: biological process (depicted in blue), cellular component (depicted in purple), and molecular function (depicted in pink). Here, each horizontal bar gives the count of genes, among the 811 human genes associated with UV exposure, linked to the respective GO term, with statistical significance (p ≤ 0.01).

In terms of biological process ontology, it can be seen that GO terms related to transcriptional processes are enriched in the set of 811 human genes (**Figure 5**), and this enrichment of transcriptional processes spans both positive and negative regulation of DNA-templated transcription [57,58]. Further, the observed GO term enrichment aligns with the established pivotal role played by transcriptional regulation in cellular responses to UV radiation [59,60]. Moreover, published literature [61] has established the association between the regulation of apoptotic processes and UV radiation exposure, and the GO enrichment analysis is in alignment with previous observations (**Figure 5**). In terms of cellular component ontology, GO terms related to intracellular membrane-bounded organelles, the nucleus, and the extracellular matrix enriched with collagen, are enriched in the set of 811 human genes (**Figure 5**), and such an association with UV radiation exposure is supported by published literature [62]. Also, as many transcriptional processes are enriched, it is logical to observe the nucleus being enriched. Further, UV radiation impacts cytoskeletal dynamics, membrane-bound organelles and focal adhesions (**Figure 5**), and this signifies the extensive influence of UV radiation exposure on cellular architecture and molecular organization. In terms of molecular function ontology, UV radiation influences distinct molecular interactions and processes which are known to shape cellular responses to environmental cues (**Figure 5**). Specifically, enriched GO terms include RNA binding and DNA binding (**Figure 5**), processes which are pivotal for transcriptional regulation [63–65]. Further, UV radiation influences protein homodimerization, receptor-ligand interactions, and cytokine activities (**Figure 5**), which suggest impact of UV radiation exposure on cellular signaling and communication pathways [65–67]. In sum, the GO analysis reveals valuable insights into the transcriptional regulatory landscape induced by UV radiation exposure. Therefore, a better understanding of the intricate interplay between UV radiation and cellular processes will be crucial for elucidating the molecular mechanisms underlying UV radiation induced responses.

Subsequently, in order to reveal the pathways significantly associated with UV radiation exposure, pathway enrichment analysis was performed to gain mechanistic insights from the set of 811 differentially expressed human genes. **Figure 6a** displays a bubble plot of the 53 enriched pathways, and it can be observed that UV radiation exposure induces a number of cellular signaling pathways. This includes the IL-17 signaling pathway which shows the highest gene overlap score, and ‘Pathways in cancer’ which is the most significant among the set of enriched pathways, and also has the maximum number of associated genes induced by UV radiation (**Figure 6a**; **Supplementary Table S7**). Of note, the IL-17 pathway plays crucial role in immune response and inflammatory reaction, while Pathways in cancer includes major signaling pathways involved in cancer development. These two pathways might interact in response to UV damage, which in turn can likely influence cellular responses such as DNA repair mechanisms and the initiation of carcinogenesis [68]. Therefore, understanding such intricate interactions is crucial for developing strategies to mitigate UV-induced damage and prevent UV-related disorders, including skin cancer [57], which remains a significant global public health concern. Further, 8 signaling pathways (out of the 53 enriched pathways) are categorized as ‘Environmental Information Processing’ in the KEGG BRITE classification at Level 1 (**Supplementary Table S7**), and these signaling pathways are also related to aging and cancer [69,70]. Furthermore, a majority of the pathways (37 out of 53) enriched within the set of 811 human genes are linked to ‘Human Diseases’ according to the KEGG BRITE classification at Level 1 (**Supplementary Table S7**). Interestingly, these 37 pathways include many viral and bacterial infection pathways. This observation can plausibly be explained by the inherent involvement of the immune system in viral and bacterial infection, a circumstance analogous to many established effects (e.g. cancer) induced by UV radiation exposure [71,72]. Furthermore, **Figure 6b** displays the pathway similarity network comprising of the 53 KEGG pathways enriched in the set of 811 human genes, showcasing the degree of similarity among the enriched pathways.

**Figure 6:**
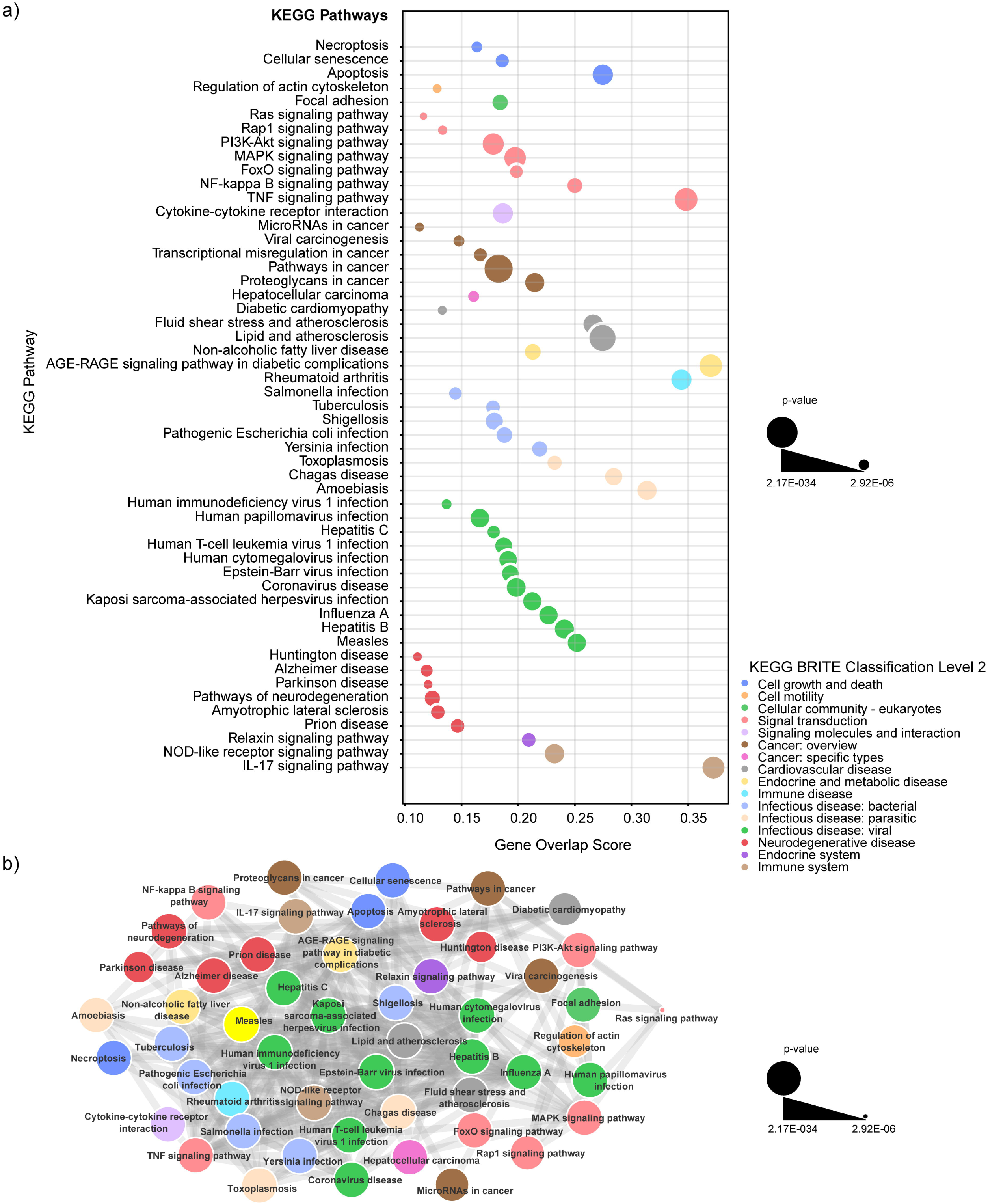
Pathway enrichment analysis of the 811 human genes induced by UV radiation exposure. **(a)** Bubble plot visualization of the 53 significantly enriched KEGG pathways in the input gene set, with bubble size reflecting significance (p-value) and placement of the bubble along the horizontal axis indicating gene overlap score (i.e., the level of overlap between input gene set and the pathway genes). Bubble colors correspond to KEGG BRITE classification at Level 2. **(b)** Pathway similarity network depicting relationships among the 53 enriched KEGG pathways in the input gene set. The similarity between any pair of pathways is quantified using the Jaccard index between the associated gene sets. Nodes are colored based on KEGG BRITE classification at Level 2, and pathways are connected if their Jaccard index exceeds 0.2.

Thereafter, to unveil the functional interactions among the 811 human genes associated with UV exposure, we analyzed the gene set within the context of the human PPI network by employing the NetworkAnalyst 3.0 webserver [51] (**Methods**). **Figure 7a** visualizes the PPI network constructed using NetworkAnalyst webserver starting with the 811 human genes associated with UV exposure. Moreover, to better understand the skin-specific responses to UV radiation exposure, we have also constructed a skin-specific PPI network (**Figure 7b**) using NetworkAnalyst webserver by considering the subset of 648 human genes reported in skin-specific experiments (**Methods**; **Supplementary Tables S10-S11**). In the constructed PPI networks, we have also marked the communities or modules with greater than 100 nodes identified by NetworkAnalyst webserver (**Figure 7**). Note that due to the high prevalence of compiled data from experiments on skin-specific tissues or cells in UVREK, the top 10 hubs or high-degree nodes in the PPI network (**Figures 7a**) corresponding to the complete set of 811 human genes are very similar to that for the PPI network (**Figures 7b**) corresponding to the subset of 648 skin-specific genes. Based on the degree, the top 10 hubs or genes in the PPI network corresponding to the 811 human genes are RPS23 (ribosomal protein S23), RPS9 (ribosomal protein S9), RPS28 (ribosomal protein S28), TP53 (tumor protein p53), RPS5 (ribosomal protein S5), UBC (ubiquitin C), RPS20 (ribosomal protein S20), CTNNB1 (catenin beta 1), RPS25 (ribosomal protein S25), and RPSA (ribosomal protein SA) (**Figure 7**; **Supplementary Table S9**). Our observation that ribosomal proteins RPS23, RPS9, RPS28, RPS5, RPS20, RPS25, and RPSA are among the top 10 hubs in the constructed PPI network, underscores the importance of protein translation processes in UV response [73,74]. Further, TP53, a well-known tumor suppressor gene, is also among the top 10 hubs, and this observation aligns with the previous reports on its role in orchestrating cellular responses to UV-induced DNA damage [75–77]. Moreover, UBC is known to be involved in protein quality control mechanisms in maintaining cellular homeostasis post UV exposure [78,79]. Further, CTNNB1 is known for its role in Wnt signaling, which influences cellular processes like proliferation and differentiation, and additionally the gene is also known to be mutated in response to UV radiation [80,60]. In sum, the elucidation of molecular interactions through a PPI network analysis contributes to a deeper understanding of UV-induced skin damage and repair mechanisms [57,77]. The analysis of the top 10 hub genes and their interactions within PPI networks will shed light on the complex molecular responses triggered by UV exposure. Furthermore, this knowledge will help in deepening our understanding of UV-induced cellular stress responses.

**Figure 7:**
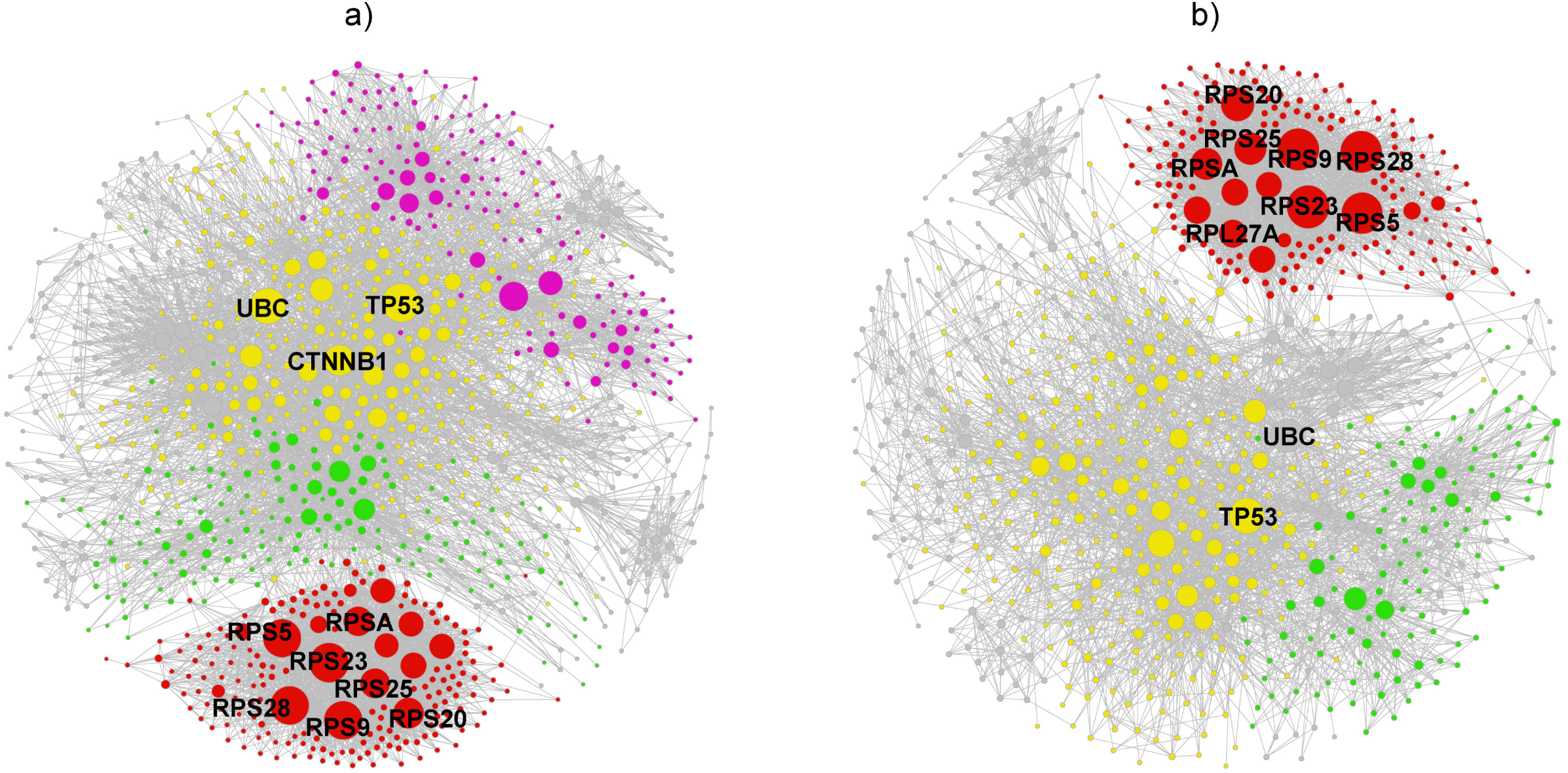
Visualization of the protein-protein interaction (PPI) network corresponding to the human genes induced by UV exposure. Visualization of the largest connected component of the first order PPI network built using: **(a)** 811 human genes. **(b)** 648 skin-specific genes. Within the PPI networks, modules of size greater than 100 are colored differently from the rest of the nodes (colored in gray). In both PPI networks, the yellow, red and green colored modules correspond to the largest, second largest and third largest, respectively, in terms of number of constituent nodes, and these large modules capture highly interconnected genes with key functional interactions induced by UV exposure. Further, the labeled nodes indicate the top 10 hub genes in the PPI networks.

## 4. Conclusions

This study is the first ever effort to create a structured database compiling information on diverse biomolecules that are induced in response to UV radiation exposure. We present UVREK database, which has compiled detailed information on 985 genes, 470 proteins, 54 metabolites, and 77 miRNAs along with corresponding metadata, through extensive manual curation from published literature. In UVREK, we have also mapped the biomolecules to the respective identifiers from standard databases, thus complying with the FAIR principles. Further, the enrichment analysis done on the gene set specific to humans revealed that the genes induced by UV exposure are involved in transcription related processes, cellular signalling and pathways related to cancer or aging.

In recent years, there is immense interest towards understanding the health impact of the different components in the human exposome. Still, the physical component of the human exposome, which includes heat, radiation and noise, remains much less characterized, hindering exploration and analysis on this front. Also, given the severity of skin disorders caused by UV radiation exposure, availability of UV radiation specific data will help in developing better diagnosis and prognosis. To this end, establishing the UVREK database with information on biomolecules induced by UV radiation exposure is an important step. Prior to our work, there were a few databases focused on radiation exposure [37,26,36] which have certain limitations. These limitations include non-availability of metadata, and restriction of data to only certain types of biomolecules. Our manually curated database, UVREK, exclusively built for UV radiation exposure addresses the limitations in the earlier databases. In other words, UVREK is a unique effort towards detailed characterization of the physical component of the exposome, by capturing the effects of UV radiation on humans and rodents.

The data in UVREK has been collected from experiments done across different organisms, specimens and measured using different methods. Hence, UVREK has the limitation of data heterogeneity. Also, the endpoints of the UV exposure were not collected, as the scope of the data collection was limited to biomolecules. This can be considered as a gap because, knowledge of the environmental factors along with their disease outcomes are crucial for risk stratification [4]. We have addressed this to an extent in this manuscript, by performing enrichment analysis which gives insights into the biological processes that might have resulted from the UV exposure. The dose data represented in our database is the total dose given to the specimen in the corresponding experimental setup and might involve multiple time points of radiation exposure. This exposure time information has not been documented explicitly in UVREK.

The compiled dataset in UVREK will be useful as a source of candidate biomarkers that can help in risk stratification and in the detection of disorders caused by UV exposure. This will be particularly useful for those exposed to artificial UV emitting equipment (e.g., water treatment, tanning beds, etc.), through formulation of proper safety guidelines. On the other hand, this data will also be useful for those who spend more time outdoors and are subject to natural UV exposure. Potential applications of the compiled data include development of better protection methods such as sunscreens. Also, this data can be a valuable resource for developing better UV sensors including UV-sensing wearables [81] that are used to detect the dose and alert people regarding their UV exposure. Lastly, it can also serve as a starting point to find new molecular signatures of UV radiation exposure.

## Data availability

The data associated with this manuscript is contained in the article or in the supplementary information or in the associated website: https://cb.imsc.res.in/uvrek/.

## Author Contributions

**Shanmuga Priya Baskaran:** Conceptualization, Data Curation, Formal Analysis, Methodology, Software, Visualization, Writing – original draft, Writing – review & editing; **Janani Ravichandran:** Conceptualization, Data Curation, Formal Analysis, Methodology; **Priya Shree:** Data Curation; **Vinayak Thengumthottathil:** Data Curation; **Bagavathy Shanmugam Karthikeyan:** Conceptualization, Formal Analysis, Supervision, Writing – original draft, Writing – review & editing; **Areejit Samal:** Conceptualization, Formal Analysis, Methodology, Supervision, Writing – original draft, Writing – review & editing.

## Acknowledgements

Areejit Samal would like to acknowledge funding from the Department of Atomic Energy (DAE), Government of India through the Apex Project to The Institute of Mathematical Sciences. The funders have no role in the study design, data collection, data analysis, manuscript preparation, or decision to publish.

## Supplementary Tables

**Table S1:** List of 2297 genes which are reported to be induced by ultraviolet (UV) radiation exposure in published studies. For each gene, we provide the corresponding Gene Symbol and NCBI Gene ID from NCBI (https://www.ncbi.nlm.nih.gov/gene/). Additionally, information on Biomolecule Type, Organism Tested, the Broad Classification of Cell/Tissue Ontology Term and the corresponding PMID are provided.

**Table S2:** List of 1228 proteins which are reported to be induced by ultraviolet (UV) radiation exposure in published studies. For each protein, we provide the corresponding UniProt ID from UniProt (https://www.uniprot.org/). Additionally, information on Biomolecule Type, Organism Tested, the Broad Classification of Cell/Tissue Ontology Term and the corresponding PMID are provided.

**Table S3:** List of 209 metabolites which are reported to be induced by ultraviolet (UV) radiation exposure in published studies. For each metabolite, we provide the corresponding CAS ID from CAS (https://www.cas.org/). Additionally, information on Biomolecule Type, Organism Tested, the Broad Classification of Cell/Tissue Ontology Term and the corresponding PMID are provided.

**Table S4:** List of 162 miRNAs which are reported to be induced by ultraviolet (UV) radiation exposure in published studies. For each miRNA, we provide the corresponding Accession from miRbase (https://www.mirbase.org/). Additionally, information on Biomolecule Type, Organism Tested, the Broad Classification of Cell/Tissue Ontology Term and the corresponding PMID are provided.

**Table S5:** The table gives the list of 811 human genes compiled in UVREK database. The PMID column lists the literature references in which the corresponding gene has been reported to be induced by UV radiation exposure.

**Table S6:** List of 30 Gene Ontology (GO) terms enriched in the set of 811 human genes induced by ultraviolet radiation (UV) exposure. The 30 GO terms are categorized into 3 ontologies namely, Biological Process, Cellular Component, and Molecular Function. For each GO term, we provide the GO Term Identifier, GO Term Name, Ontology Type, Number of Overlapping Genes from the input gene set associated with the corresponding GO term, p-value, and the Gene Symbols of the associated genes (separated by ‘;’ symbol) obtained from the Enrichr web server (https://maayanlab.cloud/Enrichr/).

**Table S7:** List of 53 pathways enriched in the 811 human genes induced by ultraviolet radiation (UV) exposure. For each pathway, we provide the corresponding KEGG ID, KEGG BRITE Classification Level 1, KEGG BRITE Classification Level 2, Number of Overlapping Genes from the input gene set associated with the corresponding pathway, Total Number of Pathway Genes, Overlap Ratio, p-value, Scaled p-value for Bubble Size and the Gene Symbols of the associated genes (separated by ‘;’ symbol) from the Enrichr web server (https://maayanlab.cloud/Enrichr/).

**Table S8:** The edge file for the largest connected first-order PPI network for the 811 human genes constructed from NetworkAnanlyst 3.0 (https://www.networkanalyst.ca/). The nodes in the network were filtered for a degree of 5. For each edge, we provide the NCBI Gene Symbol for the two nodes (Node 1 and Node 2) connected by the edge.

**Table S9:** The node file for the largest connected first-order PPI network for the 811 human genes constructed from NetworkAnanlyst 3.0 (https://www.networkanalyst.ca/). The nodes in the network were filtered for a degree of 5. For each node, we provide the NCBI Gene Symbol (represented in two columns as Id and Label), Degree and Module.

**Table S10:** The edge file for the largest connected first-order PPI network for the 648 skin-specific human genes constructed from NetworkAnanlyst 3.0 (https://www.networkanalyst.ca/). The nodes in the network were filtered for a degree of 5. For each edge, we provide the NCBI Gene Symbol for the two nodes (Node 1 and Node 2) connected by the edge.

**Table S11:** The node file for the largest connected first-order PPI network for the 648 skin-specific human genes constructed from NetworkAnanlyst 3.0 (https://www.networkanalyst.ca/). The nodes in the network were filtered for a degree of 5. For each node, we provide the NCBI Gene Symbol (represented in two columns as Id and Label), Degree and Module.

